# The response mechanism of *oreorchis patens* (Lindl.) Lindl. to cold stress

**DOI:** 10.1101/2022.07.25.501424

**Authors:** Lan Yu, Yufeng Xu, Yuyan Zhang, Meini Shao, Qing Miao, Xuhui Chen, Huixia Yang, Na Cui, Bo Qu

## Abstract

Cold stress, a major environmental factor, has an important impact on the production of landscape plants and crop yield, and its stress and resistance mechanisms have always been hot research issues. *Oreorchis pat*ens (Lindl.) Lindl., an important germplasm resource, has strong frost resistance and can resist low temperatures of -40 °C. However, the mechanism by which *O. patens* responds to cold stress remains poorly understood. Here, we examined the adaptation to the low-temperature environment of *O. patens* by changing the leaf tissue structure, while the synergistic effect of osmotic regulation, reactive oxygen scavenging and protein improved the resistance to cold stress. In addition, analysis of the photosynthetic system showed that cold stress changes the photosynthetic capacity of *O. patens* leaves to affect cold resistance. Analysis by nonparametric transcriptome sequencing revealed 2402 genes that were differentially expressed, most of which were related to resistance. Simultaneously, quantitative real-time polymerase chain reaction (qRT-PCR) analysis obtained results consistent with the transcriptome. These results indicated that *O. patens* could alter leaf structure and physiological and biochemical metabolic processes by initiating resistance-related molecular regulatory networks to improve the ability to resist cold stress. This study was the first to discuss the physiological, biochemical and molecular regulatory mechanisms of *O. patens* resistance to cold stress, which laid a foundation for revealing the biological and molecular mechanisms of overwintering of *O. patens* and breeding cold-resistant horticultural crops in northern China.

## Introduction

Cold is the main factor limiting the regional distribution of plants and an important environmental factor affecting plant growth and development. Cold stress (chilling injury and freezing injury) inhibits the growth of plants, chlorosis of leaves, wilting of plants, decrease of crop yield, and even death of plants in severe cases (Hu *et al*., 2016). Cold stress has now become an important issue that is of wide concern and urgently needs to be solved in the field of agricultural production in our country. The global annual crop loss caused by cold stress has been reported to be as high as hundreds of billions of yuan (Theochari *et al*., 2012). Therefore, exploring the physiological and biochemical response mechanism of plants under cold stress and the mechanism of cold resistance are of great significance to enhance the cold resistance of plants, increase crop yield, improve the ornamental properties of landscape plants, and breed cold-tolerant varieties of plants.

In the long-term natural selection process, plants can produce a series of adaptive responses to improve cold resistance. The morphological and anatomical structure, photosynthetic capacity, physiological and biochemical characteristics, and gene expression of plants all change under cold stress. Leaves are the main organs for photosynthesis and transpiration, and changes in leaf anatomy are the basis for plants to respond and adapt to the environment. The indicators of reactive photosynthetic capacity are the net photosynthetic rate (Pn) and intercellular carbon dioxide concentration (Cohu *et al*., 2014). The membrane system, which is sensitive to cold stress and responds to cold stress first, controls the flow of substances and signals in plants (Lyons, 1973). Several forms of damage are caused by cold stress at the cellular level, including membrane injury, metabolic imbalance of reactive oxygen species (ROS), membrane peroxidation and increase in the electrolyte permeability of the cell membrane (Liu *et al*., 2019), and inducing osmotic regulatory substance accumulation in plant cells, such as soluble sugar, soluble protein, proline, to maintain osmotic balance and reduce damage to plants (Ren *et al*., 2018). In addition, plants also activate the antioxidant enzyme system (superoxide dismutase (SOD), catalase (CAT), peroxidase (POD), etc.) to remove ROS to reduce the damage caused by reactive oxygen species to the membrane system (Li *et al*., 2019). Furthermore, cold stress can induce the accumulation of cold-responsive proteins to turn on the transduction of low temperature signals and then regulate the expression of functional genes to enhance cold resistance (Ye *et al*., 2019). With the development of high-throughput sequencing, many researchers have used RNA-Seq to study differentially expressed genes and metabolic pathways in response to cold stress to reveal the molecular mechanism of cold resistance in plants (Karki *et al*., 2013).

The germplasm resources of *Orchidaceae* in China are very abundant, but *Orchidaceae* is mainly ornamental and mostly cultivated in greenhouses due to the low temperature in the northern region. *Oreorchis patens* Lindl. is a perennial herb belonging to *Oreorchis* of *Orchidaceae*, which is widely distributed in the southwest, central, northeast and other regions of China. According to the “Chinese Materia Medica”, the pseudobulbs of *O. patens*, a highly valued medicinal, are called ice ball and alias *Cremastra appendiculata*. Studies have shown that *Cremastra* has good antibacterial properties, blocking M3 receptor, antihypertensive, antitumor, etc., activity (Tu *et al*., 2018). In addition, *O. patens* is an excellent ground cover plant with high ornamental value for landscapes.

*O. patens* with strong cold tolerance, preferring shade and humidity, can resist the low temperature of -40 °C in northeast China. *O. patens* sprouts new leaves in summer, remains evergreen in winter, and withers in the following summer. Previous studies have investigated the medicinal components, pharmacological activity, mycorrhizal fungi and reproduction of *O. patens* (Bae *et al*., 2014); however, the underlying mechanism of antihypothermia in *O. patens* has not been explored. In the limited growth period in northeast China, how can *O. patens* not only enhance cold tolerance, but also adjust growth and dormancy over time to keep its leaves green and survive in the cold winter? How does it respond adaptively?

In this study, the relationship between the morphological structure, physiological and biochemical characteristics, and cold resistance of *O. patens* were analyzed by observing the changes in the anatomical structure of *O. patens* leaves and measuring the stress resistance substances and photosynthetic system under low temperature. At the same time, nonparametric transcriptome sequencing and quantitative real-time polymerase chain reaction (qRT-PCR) were used to unravel the mechanism of cold resistance of *O. patens* leaves, providing a theoretical basis for its introduction, acclimation, artificial cultivation, and selection and identification of cold-resistant varieties of orchids.

## Results and analysis

### Changes in the anatomical structure of the leaves of *O. patens*

The leaf anatomical structure of the cross-section of *O. patens* was composed of epidermis, mesophyll and veins, which were typical of monocotyledonous leaves. The epidermis was a single layer of cells with a well-developed cuticle, the mesophyll cell structure was uniform, and the vascular bundle included the vascular bundle sheath, phloem and xylem. Cells differentiated into sclerenchyma were distributed between the vascular bundles and the upper and lower epidermis of leaves. Furthermore, *O. patens* was a C3 plant, of which the vascular bundle sheath was 1 to 2 layers of parenchyma cells and did not contain chloroplasts (Figure 1, A-F).

**Fig 1.**
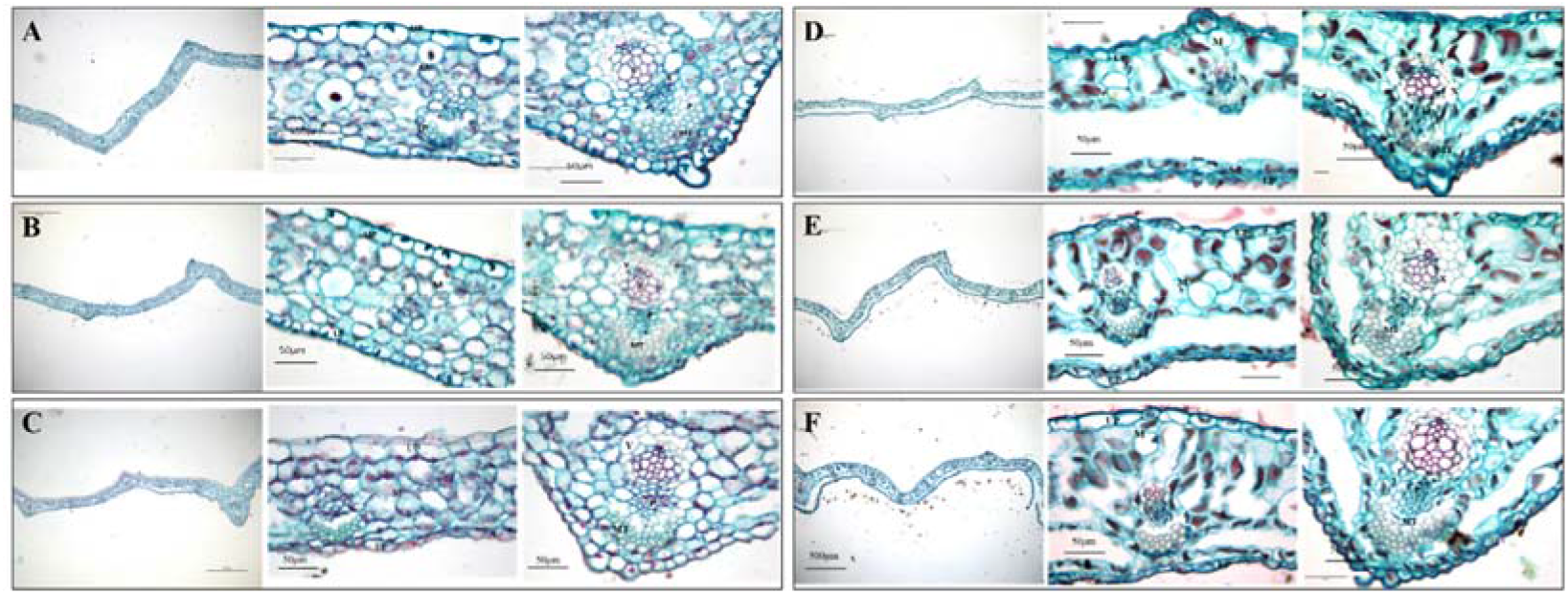
Characteristics of the leaf anatomical structure of *O. patens* at different diurnal temperatures. (A-F) Leaf cross-section structure of *O. patens* at different diurnal temperatures (day/night). Each group was the epidermis, mesophyll and leaf veins from left to right. A, 30 °C/25 °C. B, 25 °C/20 °C. C, 10 °C/5 °C. D, 5 °C/-10 °C. E, -15°C/-20°C. F, -20 °C/-30 °C. A Leica DM2500 microscope was used to observe the leaf anatomical structure of *O. patens* under different temperature treatments.

Research on the changes in the leaf anatomical structure of *O. patens* under different low-temperature treatments found that the mesophyll cells arranged neatly and tightly at higher diurnal temperatures, and as the treatment temperature gradually decreased, the degree of water loss of the mesophyll cells continued to increase, the shape gradually became irregular, and the intercellular space became increasingly larger. During this process, the thickness of the upper epidermis of the leaves increased first and then decreased, while the thickness of the lower epidermis showed a decreasing trend (Figure 2, A and B). The thicknesses of the mechanical tissue and mesophyll, numbers of mesophyll cells, xylem thickness, vessel diameter and phloem thickness first increased, then decreased and then increased with the decrease in diurnal temperature. There were significant differences among the low-temperature treatments (Fig 2, C-I). The above results indicated that cold stress changed the leaf structure of *O. patens*, which might be related to low-temperature adaptability.

**Fig 2.**
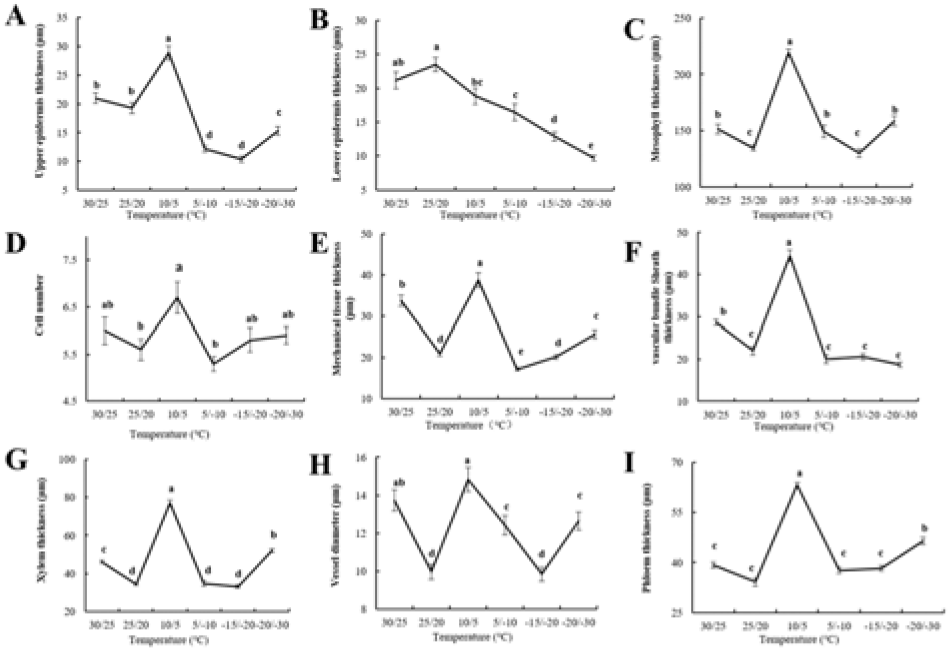
Changes in the leaf tissue structure of *O. patens* under different diurnal temperatures. The thickness of the upper epidermis (A), lower epidermis (B), thickness of mesophyll (C), numbers of mesophyll cells (D), thickness of mechanical tissue (E), thickness of vascular bundle sheath (F), xylem thickness (G), vessel diameter (H), and phloem thickness (I) were analyzed at different diurnal temperatures (day/night). Errors were standard deviations between three biological replicates (n ≥ 3). Lines annotated with different letters were significantly different according to Fisher’s least significant difference (LSD) test (*P* ≤0.05) after ANOVA.

### Effects of cold stress on the ultrastructure of mesophyll cells in *O. patens*

The ultrastructure of mesophyll cells in *O. patens* is shown in Figure 3. The ultrastructure of the chloroplasts of the mesophyll cells was observed by transmission electron microscopy (TEM). The results showed that the intact mesophyll cells had clear cell walls and membranes, clear cytoplasm and fewer flocs, in which the mostly spherical chloroplast was squeezed to the cell edge by vacuoles at higher diurnal temperatures (Fig 3, A and B). With the decrease in diurnal temperature, the gradually deformed mesophyll cell wall became unsmooth and uneven. At 10 °C/5 °C, although the chloroplast membrane began to change, most of the membranes retained their original spherical shape (Fig 3C). When the temperature dropped to 5 °C/-10 °C, the chloroplast became oblong or protruded (Fig 3D), while when the temperature dropped to -15 °C/-20 °C, some chloroplasts constricted and became amoeba-shaped (Fig 3E). Finally, the temperature dropped to -20 °C/-25 °C, and the chloroplasts disintegrated (Fig 3F). The results showed that the structure of the mesophyll was seriously affected by cold stress and even disintegrated.

**Fig 3.**
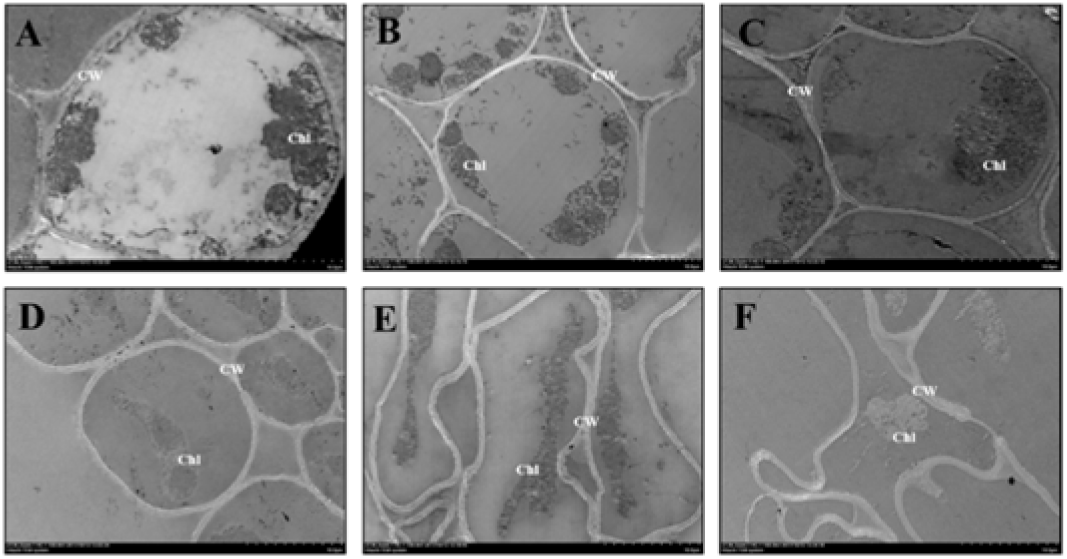
Ultrastructural analysis of the mesophyll cells. Transmission electron microscopy to observe the leaf ultrastructure of *O. patens* under different temperature treatments (day/night). (A-F) The ultrastructure of leaves at different temperatures. Chl, Chloroplast. CW, cell wall. A, 30 °C/25 °C. B, 25 °C/20 °C. C, 10 °C/5 °C. D, 5 °C/-10 °C. E, -15 °C/-20 °C. F, -20 °C/-30 °C.

### Physiological response to cold stress in *O. patens* leaves

The effects of cold stress on plants are reflected mainly in enzyme activity, membrane systems, cell water loss, etc., resulting in cell metabolism disorders and even cell death (Banerjee and Roychoudhury, 2016). During the long-term adaptation process, some plants gradually develop various cold resistance abilities, such as forming stress proteins, increasing osmotic regulatory substances, and increasing the activities of protective enzymes to improve resistance to cold stress (Lekshmi and Panigrahy, 2021). To further explore the cold-resistant characteristics of *O. patens*, we analyzed the changes in leaf-related osmotic regulators under cold stress. We found that the SOD and POD activity of leaves showed obvious unimodal curves with the change in diurnal temperature and reached a maximum at 5 °C/-10 °C (Fig 4, A and B). CAT activity increased as the diurnal temperature decreased gradually (Fig 4C). In addition, the change in MDA content also showed a unimodal curve. When the temperature was higher than 5 °C/-10 °C, MDA showed an upward trend and then gradually decreased with decreasing temperature (Fig 4D).

**Fig 4.**
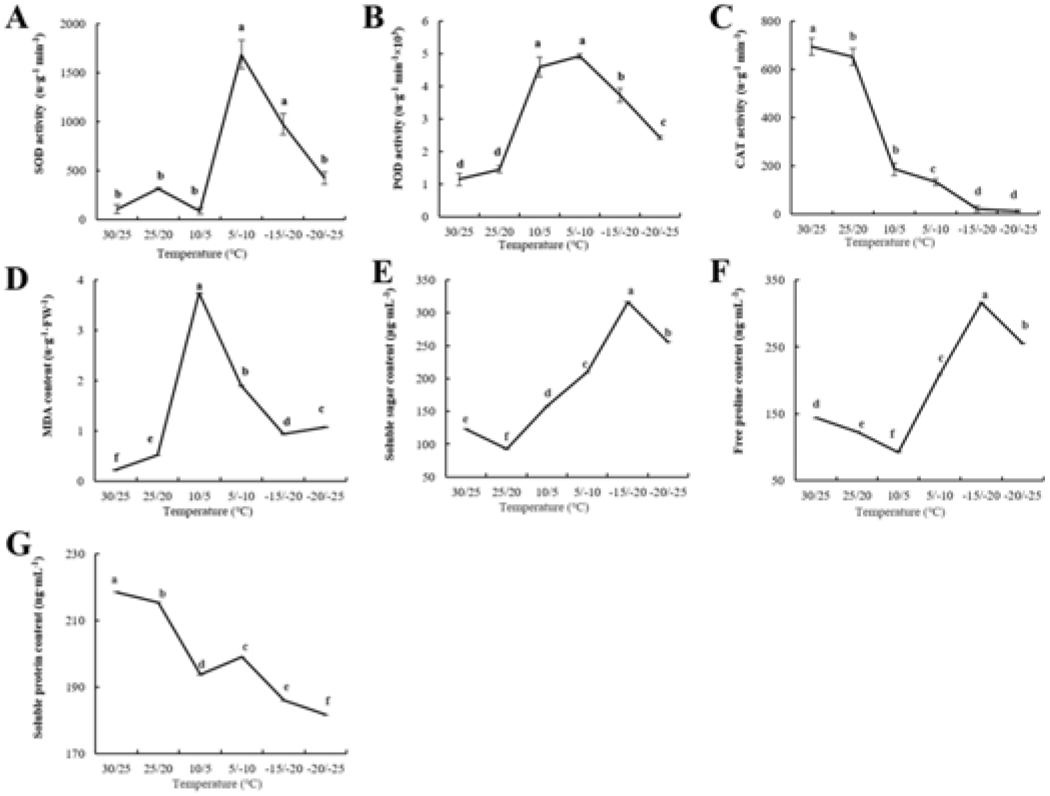
Physiological responses of leaves to cold stress. The changes in SOD activity (A), POD activity (B), CAT activity (C), MDA content (D), soluble sugar content (E), free proline content (F), and soluble protein content (G) of *O. patens* were analyzed at different diurnal temperatures (day/night). Errors were standard deviations with three biological replicates (n ≥ 3). Lines annotated with different letters were significantly different according to Fisher’s LSD test (*P* ≤ 0.05) after ANOVA.

Cold tolerance has been reported to be linked to adaptation or freezing tolerance in plants. Biomolecules such as soluble sugars, proline and various low-molecular-weight solutes are reported to protect plants from cold stress (Ruelland *et al*. 2009). In this study, the content of soluble sugar and free proline in leaves gradually increased with decreasing diurnal temperature, and the content reached a maximum at -15 °C/-20 °C (Fig 4, E and F). However, the change in soluble protein content was different from the former, which decreased significantly with the decrease in diurnal temperature (Fig 4G). These results indicated that *O. patens* could adapt to a low-temperature environment by adjusting various osmotic substances.

### Changes in photosynthesis in leaves of *O. patens* under cold stress

Photosynthesis is the physiological process that determines plant growth and yield and ultimately affects survival. However, plants exposed to cold stress exhibit changes in photosynthetic efficiency (Banerjee and Roychoudhury, 2018). In this study, we found that the chlorophyll content, transpiration rate, stomatal conductance and net photosynthetic rate of *O. patens* decreased significantly with decreasing diurnal temperature (Fig 5). When the diurnal temperature was higher than 10 °C/5 °C, the chlorophyll content did not change significantly but decreased significantly below this temperature (Fig 5A). When the diurnal temperature dropped from 35 °C/25 °C to 25 °C/20 °C, the net photosynthetic rate of the leaves increased significantly. When the diurnal temperature dropped to 10 °C/5 °C, the net photosynthetic rate decreased to the minimum, and the net photosynthetic rate approached zero as the temperature dropped further (Fig 5B). The changes in stomatal opening were shown to be responsible for the altered photosynthetic efficiency (Paul *et al*., 2008). To explore the reasons for the change in photosynthetic rate, we measured the stomatal conductance and transpiration rate of leaves to show that the variation trends of leaf stomatal conductance and transpiration rate tended to be consistent. When the diurnal temperature was higher than 10 °C/5 °C, the leaf stomatal conductance and transpiration rate first decreased, then increased, and then showed a sharp downward trend with the decrease in temperature (Figure 5, C-E). In addition, the intercellular CO_2_ concentration in the leaves showed a bimodal curve with the change in diurnal temperature. When the diurnal temperature dropped to 10 °C/5 °C, the intercellular CO_2_ concentration reached the first peak and then decreased. At -15 °C/-20 °C, the intercellular CO_2_ concentration increased and reached the second peak (Fig 5D). These results indicated that cold stress might change the net photosynthetic rate by affecting the chlorophyll content, transpiration rate and stomatal conductance in leaves of *O. patens*.

**Fig 5.**
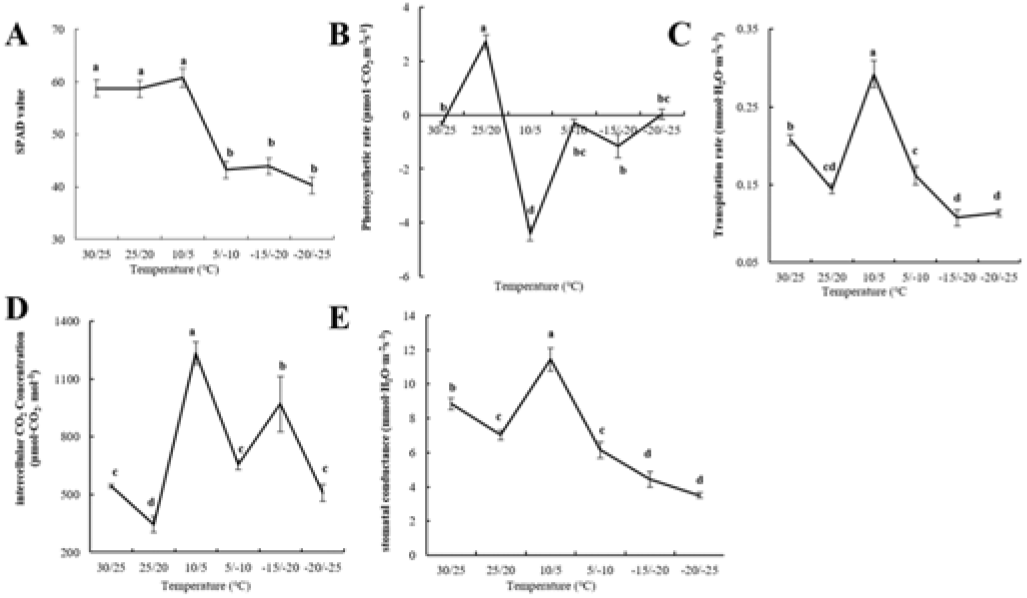
Changes in photosynthesis in leaves of *O. patens* under cold stress. The changes in soil plant analysis development (SPAD) (A), photosynthetic rate (B), transpiration rate (C), intercellular CO_2_ concentration (D), and stomatal conductance (E) of *O. patens* were analyzed at different diurnal temperatures (day/night). Errors were standard deviations with three biological replicates (n ≥ 3). Lines annotated with different letters were significantly different according to Fisher’s LSD test (*P* ≤ 0.05) after ANOVA.

### Analysis of transcriptome changes in overwintering *O. patens*

The key to understanding the plant cold response lies in identifying the possible molecular mechanisms of temperature sensing and signaling (Dai *et al*., 2007). To further study the resistance mechanism of *O. patens* to low temperature, transcriptome analysis of *O. patens* at the overwintering stage was performed. The Gene Ontology (GO) enrichment results of the nonparameter transcriptome in leaves of *O. patens* showed that there were a total of 43 GO pathways enriched for differentially expressed genes in leaves of *O. patens* compared with the control group. The most prominent GO pathways in the molecular function classification were structural molecule activity and structural constituent of ribosome. Cytoplasm and intracellular organelle were significantly enriched pathways in the classification of cellular components, indicating that the main functional genes and their encoded proteins and noncoding RNAs were located mainly in the cytoplasm of *O. patens* during overwintering (Figure 6A, Table 1). To gain an in-depth understanding of the changes in signal transduction and metabolic pathways in the overwintering period of *O. patens*, we performed Kyoto Encyclopedia of Genes/Genomes (KEGG) enrichment analysis. A total of 180 differentially expressed genes were enriched in 49 metabolic pathways, of which 34 articles were enriched in metabolic processes. The enrichment results showed that the overwintering of *O. patens* was mediated mainly by the increase in the cutin, suberine and wax signaling pathways and the decrease in the peroxisome signaling pathway (Figure 6B, Table 2). We screened the selected differentially expressed genes (DEGs) and obtained 2,402 DEGs, including 766 downregulated genes and 1,636 upregulated genes (Figure 6C, Table 3). The functional predictions were divided mainly into genes related to hormone synthesis and response, cold shock proteins, C-repeat binding transcription factor (CBF), osmotic regulation substance synthesis, etc. In summary, the transcriptome results showed that *O. patens* might respond to cold stress by regulating the hormone content, increase or decrease in osmotic substances, and cold shock proteins.

**Fig 6.**
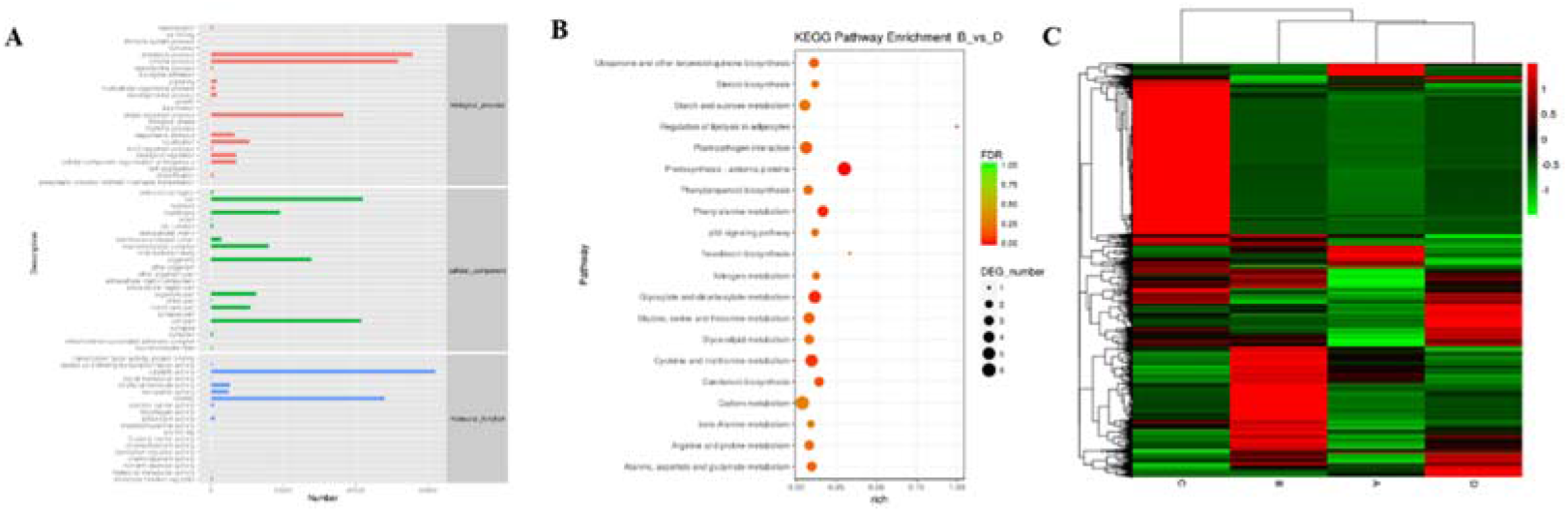
Transcriptome analysis of the overwintering leaves of *O. patens*. (A) Results of GO classification and annotation by nonparametric transcriptome. (B) KEGG enrichment analysis of differentially expressed genes. (C) Heatmap analysis of differential gene expression.

**Table 1.**
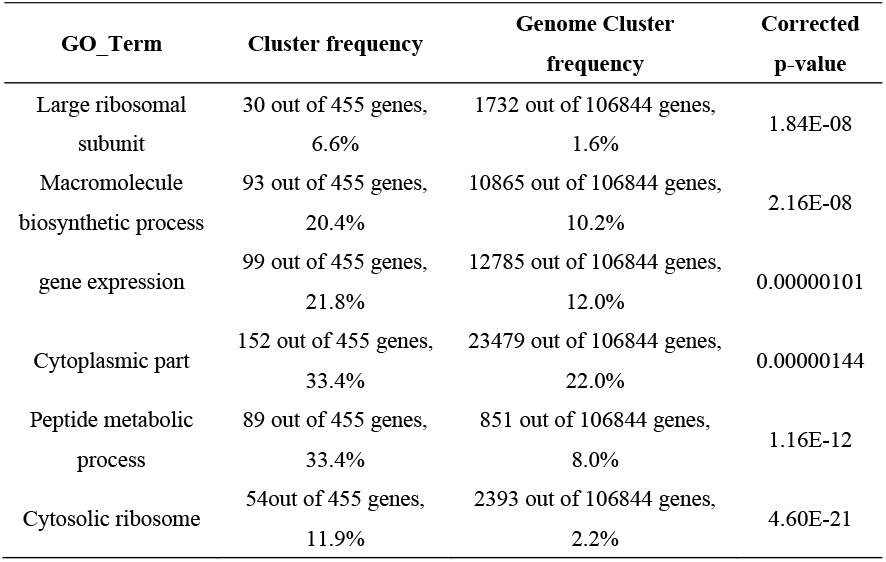
Enrichment analysis of functional significance on the Gene Ontology

**Table 2.** KEGG pathway enrichment analysis of differentially expressed genes

| Pathway_ | Pathway_ ID | DEC_ number | Total_ number | pvalue |
| --- | --- | --- | --- | --- |
| Cutin, suberine and wax biosynthesis | ko00073 | 3 | 7 | 2.60967E-05 |
| Longevity regulating pathway - worm | ko04212 | 3 | 39 | 0.005524654 |
| Renin-angiotensin system | ko4614 | 1 | 2 | 0.01859884 |
| Peroxisome | ko4614 | 3 | 63 | 0.01968867 |
| alpha-Linolenic acid metabolism | ko00592 | 2 | 25 | 0.02236359 |
| Tyrosine metabolism | ko00350 | 2 | 26 | 0.02408463 |
| Oxidative phosphorylation | ko00190 | 4 | 120 | 0.02498221 |
| Amino sugar and nucleotide sugar metabolism | ko00520 | 3 | 73 | 0.03015742 |

**Table 3.**
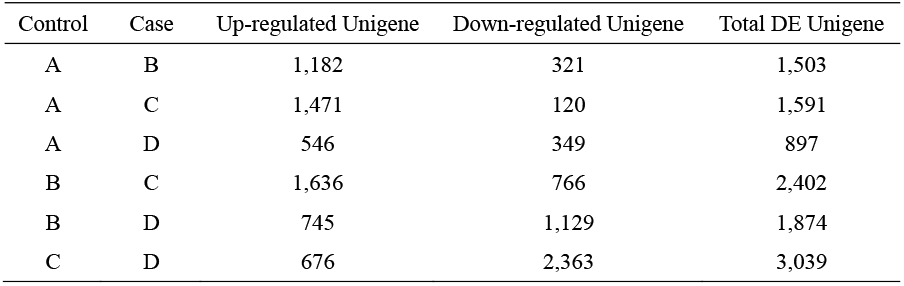
Statistical analysis of differentially expressed genes

### Cloning of the *Actin* fragment in *O. patens*

At present, there are relatively few studies on *O. patens*, and its *Actin* gene has not yet been cloned. Thus, we downloaded the *Actin* nucleotide sequences of *Orchidaceae*, namely, *Phalaenopsis Aphrodite, Doritaenopsis hybrid, Cymbidium ensifolium, Oncidium hybridum*, and *Sedirea japonica*, from the National Center for Biotechnology Information (NCBI) database (https://www.ncbi.nlm.nih.gov/). We performed homology comparisons using DNAMAN to identify highly conserved segments (Fig 7A). A pair of primers were designed by Primer 5.0 biological software; the forward primer was 5’-AGCAACTGGGATGATATGGAAAA-3’, and the reverse primer was 5’-GCTTGAATGGCAACATACATGG-3’, which were synthesized by Jinweizhi Biotechnology Co., Ltd. To amplify the *Actin* sequence of *O. patens*. The designed primers were used for gel electrophoresis analysis, which showed that the amplified bands were between 100-200 bp (Figure 7B). Then, the band was analyzed by gel recovery and sequencing by Jinweizhi Biotechnology Co., Ltd. The homology obtained sequences were compared with other *Orchidaceae Actins*, which found that the homology reached 90.8% (Fig 7C). These results indicated that the *Actin* fragment was cloned successfully, and the subsequent experiments could be used for qRT-PCR analysis.

**Fig 7.**
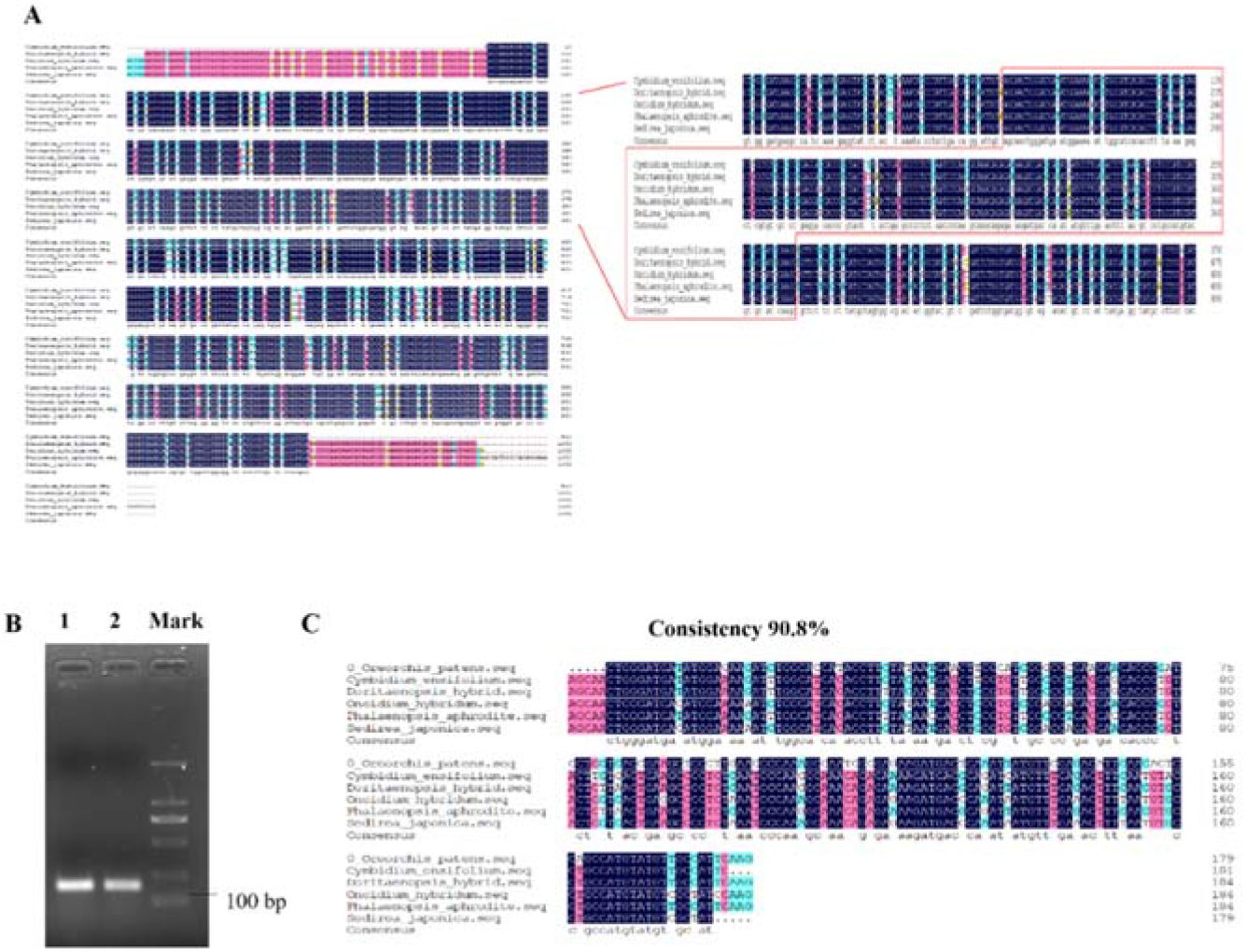
Cloning the *Actin* gene of *O. patens*. (A) Homologous alignment analysis of *Actin* sequences in different *Orchidaceae* plants. DNAMAN was used for homology alignment analysis. The homologous alignments of *Orchidaceae* were *Phalaenopsis Aphrodite, Doritaenopsis hybrid, Cymbidium ensifolium, Oncidium hybridum*, and *Sedirea japonica*. (B) Electrophoresis detection of *Actin* in *Oreorchis*. 1-2, PCR production of *Actin*. M, DNA Marker of DL2000. (C) Similarity analysis of the *Actin* sequence of *Oreorchis* with other *Orchidaceae*. DNAMAN was used for the analysis.

### Verification of differentially expressed genes under cold stress by qRT-PCR

To verify the transcriptome data, *O. patens* was subjected to cold stress treatment at 4 °C for 24 h, and samples were taken for qRT-PCR. We screened 3 upregulated genes, namely, *c433773-g1, c435999-g1* and *c446826-g1*. Transcriptome prediction analysis showed that these 3 genes were annotated as proline receptor protein kinase 3 (PERK3), serine/threonine interacting with calcineurin B-like (CBL) protein kinase, and salicylic acid binding protein 2 (SABP2) (Zhu *et al*., 2015). Three genes were significantly downregulated, namely, *c182937-g1, c466636-g1* and *c472004-g1*. These genes were annotated as genes related to abscisic acid production, maize transcription factor BTF3, and cold shock protein.

Through transcriptome prediction and a literature review, we learned that these 6 genes were all involved in the plant cold stress response. Studies have found that PERK can act as a sensor. When plants are subjected to biotic and abiotic stresses, PERK monitors changes in plant cell walls and activates relevant cellular responses to respond to stress (Doblin *et al*., 2014). Salicylic acid, a phenolic compound, can promote the growth and development of plants. External application of salicylic acid can improve the tolerance of plants to abiotic stresses, such as cold stress, drought, and salt stress (Siboza *et al*., 2014; Guan *et al*., 2019). A study on lemon fruit found that external application of salicylic acid can improve the cold tolerance of lemons (Siboza *et al*., 2014). SABP2 can play an active regulatory role in the plant stress response by increasing endogenous SA levels (Guan *et al*., 2019). Studies have shown that CBL can respond to biotic and abiotic stresses through the CBL-CIPK pathway, including cold stress (Ma *et al*., 2019; Ma *et al*., 2020). Abscisic acid (ABA) is a crucial phytohormone that has been reported to regulate many important physiological and biochemical processes in plants under abiotic stress. Studies on *Cynodon dactylon* find that ABA can regulate the response of plants to cold stress by regulating electrolyte leakage, MDA, H_2_O_2_, etc (Huang *et al*., 2017). Basic transcription Factor 3 (BTF3) is the β-subunit of the nascent polypeptide-related complex, which is responsible for the transcription initiation of RNA polymerase II and participates in cell apoptosis, translation initiation regulation, growth, development and other processes. Studies have shown that *TaBTF3* silencing reduces the survival rate, free proline content and relative water content of wheat plants but increases the relative electrical conductivity and water loss rate, indicating that *TaBTF3* silencing reduces wheat tolerance to cold and drought stress (Kang *et al*., 2013). In plants, cold shock proteins can enable plants to acquire freezing tolerance (Behl *et al*., 2020). In a study of *Arabidopsis*, CSDP1 and CSDP2 were found to reduce freezing injury and increase the cold resistance of *Arabidopsis* (Park *et al*., 2009).

By analyzing the expression of these genes, we found that the transcript levels of *c433773-g1, c435999-g1*, and *c446826-g1* were significantly increased (Fig 8A), and the transcript levels of *c182937-g1, c466636-*g1 and *c472004-g1* were significantly decreased at 4 °C (Fig 8B), consistent with the transcriptomic data results. These results indicated that the resistance of *O. patens* to cold stress might affect the related physiological functions by regulating the gene expression related to resistance and improve the ability to resist low temperature to survive smoothly in the winter.

**Figure 8.**
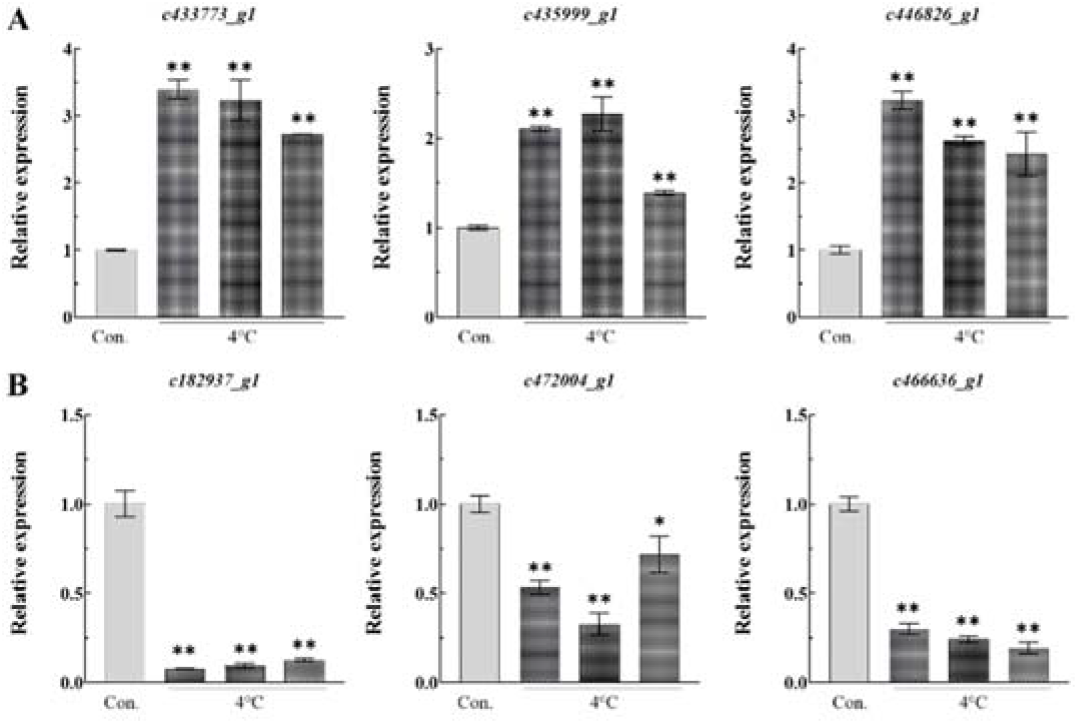
Validation analysis by qRT-PCR. The plants of *O. patens* were treated with 4 °C, and the relative transcript levels of significantly different genes of upregulated genes (A) and downregulated genes (B) for transcriptome analysis were analyzed. *c433773-g1, c435999-g1*, and *c446826-g1* were annotated as proline receptor protein kinase PERK3, serine/threonine interacting with CBL protein kinase, and salicylic acid binding protein SABP2. *c182937-g1*, c466636-g1, and *c472004-g1* were annotated as genes related to abscisic acid production, maize transcription factor BTF3, and cold shock protein. Control (Con.), 4 °C treated (4 °C). All transcript levels were determined by qRT-PCR and normalized to *Actin* transcript levels. Errors were standard deviations with three biological replicates (n ≥ 3). Differences between means were analyzed for significance using Fisher’s LSD test, \**P* < 0.05, \*\**P* < 0.01. ns, no significant difference.

### Discussion

Temperature is the most basic determinant and key environmental factor that affects plant growth and development and even causes plant death. Among these factors, low temperature can change the morphological and anatomical structure as well as a series of physiological and biochemical processes in plants.

Leaves are an important organ of plants exposed to the environment. The morphological structure is closely related to the surrounding environment, with plasticity and an obvious response to environmental changes (Tian *et al*., 2020). Low temperature is the most important ecological factor affecting the growth and development of plant leaves. Plant leaves can adapt to cold stress and improve their resistance by changing their morphological and structural characteristics (Zhang *et al*., 2016). The epidermis of the leaf is a vital protective tissue, so the thicker the upper and lower epidermises of the leaves are, the stronger the water retention capacity and the more able the plant is to resist the damage caused by low temperature (Chen *et al*., 2018). The research by Liu (2004) on the cold resistance of 7 species of *Annona squamosal* Linn. found that the greater the protuberance of the leaf veins, the more water loss can be compensated by the leaf vein vascular tissue to accelerate the material transport capacity. These strategies enhance its cold resistance (Liu *et al*., 2004). The diameter of xylem vessels, the width of phloem tissue, and the number of sclerenchyma in leaves in nine of *Stevia rebaudiana* are increased under cold stress (Hajihashemi *et al*., 2018). Low temperature has been reported to lead to a significant increase in the number of phloem cells in leaf veins, thereby increasing sugar export from leaves to other parts of the plant (Stewart *et al*., 2016). Furthermore, water transport is related mainly to the structure of the xylem vessels, while narrow conduits offer high resistance during transport but are less prone to cavitation, which can provide safety during winter freezing (Psaras and Sofroniou, 2004). The results of this study showed that the thickening of the lower epidermis, the thickness of mechanical tissue, mesophyll, phloem and xylem, and the diameter of the vessel in *O. patens* with the decrease in diurnal temperature showed a trend of first increasing, then decreasing and then increasing, which was the adaptive response of *O. patens* to cold stress.

Cold stress may cause plasmolysis and enlargement of the cell gap, cause damage to the chloroplast membrane, cause changes in the membrane components and membrane proteins of the thylakoid membrane, and destroy the ultrastructure of the thylakoid inner membrane (Popov and Astakhova, 2021). A similar phenomenon was also observed in our study, in which the chloroplasts of the leaves were gradually dissolved, the numbers were decreased, and the cell wall was deformed with decreasing temperature. However, from the perspective of the whole cooling process, the chloroplast could still maintain integrity when the temperature dropped to -20 °C, which indicated that *O. patens* had strong cold tolerance (Hao *et al*., 2017).

Cold stress leads to a decrease in plant photosynthetic activity, an increase in cell membrane permeability, and a series of changes in physiological and biochemical properties, which ultimately inhibit plant photosynthesis (Cohu *et al*., 2014). Some studies have shown that cold stress reduces the photosynthetic rate and pigment content to inhibit photosynthesis (Strand *et al*., 1997). Cold stress inhibits the biosynthesis of 5-aminolevulinic acid in plant leaves to reduce chlorophyll synthesis (Mohanty *et al*., 2006). The reduction of photosynthetic pigments reduces the level of light absorption and subsequently the level of excess excitation energy as an alternative to increasing the level of energy dissipation, which reduces photosystem II complex (PSII) efficiency (Demmig-Adams and Adams, 2006). Demmig-Adams (2006) found that mesophytes tend to promote photosynthetic capacity and increase chlorophyll levels in winter, while several evergreens and conifers tend to enhance photoprotection and retain chlorophyll, which relies more on higher levels of energy dissipation (Demmig-Adams *et al*., 2006). This study showed that the chlorophyll content in the leaves of *O. patens* did not change significantly when the diurnal temperature was higher than 10 °C/5 °C, but decreased significantly when the temperature was lower than 5 °C/-10 °C.

Under low-temperature conditions, the photosynthetic rate in the leaves of *O. patens* was significantly reduced, which was an adaptive response to cold stress. Studies have shown that photoinhibition is characterized by reduced photosynthetic CO_2_ absorption. An increase in the concentration of intercellular CO_2_ favors photosynthesis, whereas the opposite indicates that the photosynthetic organ is inhibited (Adams *et al*., 2013). Zhu (2018) found that an increase in intercellular CO_2_ concentration improves the photosynthetic electron transfer efficiency of wheat plants under cold stress (Zhu *et al*., 2018). The results of this study showed that the intercellular CO_2_ concentration in the leaves of *O. patens* was significantly reduced under cold stress, which resulted in a decrease in the absorption of photosynthetic CO_2_, thereby inhibiting photosynthesis. Tian (2020) found that the transpiration rate and stomatal conductance of varieties with strong cold resistance are lower than the transpiration rate and stomatal conductance of varieties with weak cold resistance under cold stress (Tian *et al*., 2020). In this study, we found that the main reason for the reduction in the photosynthetic rate was that chloroplast development and chlorophyll synthesis were inhibited by cold stress. In addition, the stomatas in the leaves were closed, and the intercellular CO_2_ concentration was decreased, which resulted in a reduction in the transpiration rate under cold stress.

To adapt to low temperature, the water-holding capacity of cells can be enhanced by changing hydrophilic substances, such as soluble proteins and soluble sugars in plant cells, to improve the water-absorbing and water-holding capacity of cells and reduce the extravasation of water from cells to avoid dehydration and coagulation of protoplasm, which enhances the cold tolerance of plants (Ritonga *et al*., 2020). The content of soluble protein and soluble sugar are positively correlated with cold resistance, which has been proven in a variety of plants. Research has shown that the soluble sugar content in different grape varieties increases under low-temperature conditions, but the soluble sugar accumulation of varieties with strong cold tolerance is higher than the soluble sugar content of sensitive varieties (Grant *et al*., 2009). Low temperature can induce a high soluble protein content in barley leaves, but there are differences in protein profiles and densities between different varieties and temperature treatments, and winter barley varieties respond more rapidly to low temperature than spring barley varieties (Karimzadeh *et al*., 2005). In our study, the content of soluble protein and soluble sugar increased significantly with decreasing temperature (*P* < 0.05), which indicated that *O. patens* could improve its self-resistance to cold by maintaining a high content of soluble protein and soluble sugar to reduce damage. Meanwhile, studies have shown that cold stress could result in a large amount of free proline accumulation, and the mass fraction of intracellular free proline was positively correlated with plant cold resistance. However, some studies have pointed out that the accumulation of proline is only to adapt to the low temperature environment (Hanson and Hitz, 1982). In this study, the proline content in the leaves of *O. patens* decreased at the early stage of cooling, but the proline content gradually increased with a further decrease in temperature, indicating that the leaf cells of *O. patens* might decompose some proteins into amino acids to reduce the cell osmotic potential and improve cellular water retention.

Protective enzymes SOD, POD and CAT are important for scavenging free radicals in plants. Under normal circumstances, the production of free radicals in plant cells and the scavenging system composed of oxidase are a dynamic balance that controls ROS in plant cells at a low level, thereby protecting cells from damage (Tang *et al*., 2019). Research on the molecular regulation mechanism of tomato finds that overexpression of the cold resistance gene *LeCOR413PM2* increases the activities of SOD, POD and CAT to reduce the accumulation of ROS, which increases the cold resistance of tomato (Zhang *et al*., 2021). In this study, we found that the SOD activity of the leaves was significantly correlated with the cold tolerance of the plant, indicating that *O. patens* could improve the cold resistance by changing the SOD activity. In addition, the activity of POD increased with decreasing temperature, but there was little change compared with the change in other plants, indicating that POD did not play a major role in cold tolerance. The CAT activity increased with decreasing temperature and was always higher than the CAT activity of other plants, indicating that the CAT activity of *O. patens* was significantly correlated with its cold tolerance. In addition, the scavenging ability of ROS also contributed greatly to cold resistance. The change in MDA content can reflect the degree of membrane lipid peroxidation, thus reflecting the degree of plant damage under environmental stress (Fan *et al*., 2012). Numerous studies have shown that cold stress increases the permeability of plant cell membranes and significantly increases the MDA content. In this study, the MDA content varied with decreasing temperature. When the temperature dropped to 5 °C, the cells were sensitive, and the degree of membrane peroxidation was high, which caused cell damage. However, when the temperature dropped below 5 °C, the degree of peroxidation of the cell membrane slowed down, indicating that *O. patens* had a strong ability to resist cold stress.

Transcriptome sequencing (RNA-seq) based on next-generation sequencing technology can align hundreds of millions of sequencing results to a reference genome (parametric transcriptome). Transcriptome-level analysis also enables the discovery of novel transcripts that aid in refining reference genome annotations. For species without a reference genome, the results of transcriptome sequencing can obtain a complete mRNA set through de novo splicing (parameter-free transcriptome) and can also perform transcript level analysis, which enables the whole transcriptome analysis of nonmodel species. In this study, no-parameter transcriptome sequencing was performed on leaves of *O. patens* treated with low temperature. The results of differential gene screening (logFC ≥ 2, *P* ≤ 0.05) showed that there were a total of 2402 differential genes, of which a total of 1636 genes were upregulated, but 766 were downregulated. The results of GO enrichment and KEGG comparison showed that there were 43 GO pathways and 50 KEGG pathways with the most significant expression differences, which laid a biological foundation for further exploring the biological and molecular mechanism of overwintering in *O. patens*. The results of differential gene expression by qRT-PCR screening were consistent with the transcriptome results, indicating that *O. patens* might improve cold tolerance through the hormone pathway, inducing the expression of cold shock proteins and cold-related transcription factors, but the specific regulatory mechanism needs to be further analyzed, which also laid the foundation for further exploration of the molecular regulatory mechanism of cold tolerance in *O. patens*.

Based on the analysis of the effects of cold stress on the physiological and biochemical and transcriptome changes in *O. patens*, we speculated that *O. patens* mainly regulated the related hormone pathways, such as the abscisic acid pathway, by initiating the expression of genes related to hormone synthesis and response to improve cold resistance. In addition, low temperature might possibly stimulate the expression of resistance-related protein genes, such as *CBL*, to improve the cold resistance of *O. patens*. However, its specific regulatory mechanism is still unknown, so we will further explore these two regulatory mechanisms in future research.

## Materials and Methods

### Plant materials and growth conditions

The *Oreorchis patens* (Lindl.) Lindl was used in this study. In the autumn of 2020, plants with consistent growth were selected, planted in cylindrical culture pots with a diameter×depth of 10 cm×10 cm, and cultivated in a light incubator (12 h light/12 h dark cycle) with a light intensity of 5500 Lux. During this period, the plants were watered one time each day to ensure that the growth environment of *Oreorchis* was humid.

Six day/night temperature gradients were set up according to the characteristics of the phenophase of *O. patens*, and the processing sequences were 30 °C/25 °C, 25 °C/20 °C, 10 °C/5 °C, 5 °C/-10 °C, -15 °C/-20 °C, and -20 °C/-30 °C. After culturing for 10 d in each temperature treatment process, the photosynthetic system indices of the leaves were measured, and then the leaves were collected in 3 groups. The first group was fixed with FAA (10% formalin, 5% acetic acid, 50% ethanol) for microscopic observation, the second group was fixed with glutaraldehyde for ultrastructure observation, and the third group was stored at -20 °C for the determination of osmotic regulators, membrane lipid peroxidation products, enzymatic activity, and transcriptome sequencing.

### Organizational structure observation

The leaf microstructure of *O. patens* under different low-temperature conditions was observed by paraffin sectioning. A piece of 5 mm×3 mm was cut from the middle of the leaf, which was fixed in FAA, dehydrated by alcohol gradient dehydration, transparent, immersed in wax, embedded, sectioned by paraffin microtome (thickness 8∼10 μm), stained with safranin, sealed with Canadian gum, finally observed and photographed under a Leica DM2500 microscope (Leica Biosystems, Shanghai, China). At the same time, the thickness of the upper and lower epidermis, mesophyll, mechanical tissue, vascular bundle sheath, xylem, phloem, the numbers of mesophyll cells, and the diameter of the vessel were measured. Three sections were observed for each temperature treatment, and 10 fields of view were taken for each section with the average value.

The ultrastructure of the chloroplasts in mesophyll cells of *O. patens* under different low-temperature conditions was observed by transmission electron microscopy (TEM). Another small piece of leaf was taken, fixed with glutaraldehyde, dehydrated, soaked, sliced, stained, observed and photographed with a TEM-1200EX electron microscope.

### Photosynthetic analysis

The SPAD-502plus instrument (Konica Minolta Company, Shanghai, China) was used to measure the chlorophyll content in the leaves of *O. patens*. An LI-6400XT portable photosynthesis instrument (LI-COR, Lincoln, NE, USA) was used to measure Pn (net photosynthetic rate), Tr (transpiration rate), Ci (changing intercellular CO_2_ concentration), and Gs (leaf level stomatal conductance) in the leaves of *O. patens*. Five plants were tested for each treatment, and each plant was tested three times. Measurements were performed at approximately 10:00 am every day, and data were recorded.

### Determination of malondialdehyde (MDA) and osmotic adjustment substances

MDA was measured by thiobarbituric acid (TBA). Soluble protein content was measured by coomassie brilliant blue G250, soluble sugar was measured by anthrone colorimetric, and free proline (Pro) content was measured by ninhydrin-sulfosalicylic acid.

### Assay of antioxidant enzyme activity

The activity of superoxide dismutase (SOD) was measured by nitroblue tetrazolium, peroxidase (POD) was measured by guaiacol, and catalase (CAT) was measured by spectrophotometer.

### Parametric transcriptome analysis

Parametric transcriptome analysis was performed by Shanghai Parsenor Biotechnology Co., Ltd. (Shanghai, China).

### RNA extraction and qRT-PCR

The functional leaves of the low-temperature-treated *O. patens* were taken and frozen with liquid nitrogen. Total RNA from plants was extracted using Trizol-reagent (CWBIO, Jiangsu, China). Bioshap was used to measure the RNA concentration, and cDNA was synthesized by the FastKing cDNA First-Strand Synthesis Kit (TIANGEN, Beijing, China). qRT-PCR was performed using a SYBR Green PCR Master Mix kit on a CFX96 Touch Real-Time PCR Detection System (Bio-Rad, Hercules, CA, USA). The reaction system (10 μL) included 2×SuperReal PreMix Plus 4.5 μL, cDNA 1 μL, upstream primer 0.5 μL, downstream primer 0.5 μL, and finally made up to 10 μL with double-distilled (dd)H_2_O. The reaction program consisted of 40 cycles of 95 °C for 15 min, 95 °C for 10 s, and 60 °C for 32 s, and the dissolution curve program consisted of 65 °C for 5 s and 95 °C for 5 min.

### Data processing

All experiments in this study were performed with at least three repetitions. Experimental data were processed using Microsoft Excel 2010 and GraphPad Prism 6 for graphing. The significance of differences was determined by analysis of variance (ANOVA) or Student’s *t-*test using IBM SPSS 20 software. Duncan’s test for multiple comparisons for significant differences (*P*<0.05) was performed. The data in the graphs are the mean ± SD (standard deviation).

## Author Contributions

LY and YX contributed to the experimental design, experimenting and result analysis and writing. YZ, MS and QM contributed to the planting and sampling and data processing. XC and HY contributed to analyze the data in experiment. NC and BQ revised the paper. All authors read and approved the final manuscript.

## Acknowledgments

We would like to thank professor Guifang He of Qinghai University for her assistance in collecting samples, and the Special Project of Orchid Survey of National Forestry and Grassland Administration for providing financial support.

